# Evolutionary dynamics of CRISPR gene drives

**DOI:** 10.1101/057281

**Authors:** Charleston Noble, Jason Olejarz, Kevin M. Esvelt, George M. Church, Martin A. Nowak

## Abstract

The alteration of wild populations has been discussed as a solution to a number of humanity’s most pressing ecological and public health concerns. Enabled by the recent revolution in genome editing, CRISPR gene drives, selfish genetic elements which can spread through populations even if they confer no advantage to their host organism, are rapidly emerging as the most promising approach. But before real-world applications are considered, it is imperative to develop a clear understanding of the outcomes of drive release in nature. Toward this aim, we mathematically study the evolutionary dynamics of CRISPR gene drives. We demonstrate that the emergence of drive-resistant alleles presents a major challenge to previously reported constructs, and we show that an alternative design which selects against resistant alleles greatly improves evolutionary stability. We discuss all results in the context of CRISPR technology and provide insights which inform the engineering of practical gene drive systems.

## Main Text

Gene drive systems are selfish genetic elements which bias their own inheritance and spread through populations in a super-Mendelian fashion (Fig. 1A). Such elements have been discussed as a means of contributing to the eradication of insect-borne diseases such as malaria, reversing herbicide and pesticide resistance in agriculture, and controlling destructive invasive species (*1–12*).Various examples of gene drive can be found in nature, including transposons (*13*), Medea elements (*14, 15*), and segregation distorters (*16–19*), but for ecological engineering purposes, endonuclease gene drive systems have received the most significant attention in the literature (*1–10, 20–22*). In general, these elements function by converting drive-heterozygotes into drive-homozygotes through a two-step process: (i) the drive construct, encoding a sequence-specific endonuclease, induces a double-strand break (DSB) at its own position on a homologous chromosome, and (ii) subsequent DSB repair by homologous recombination (HR) copies the drive into the break site. Any sequence adjacent to the endonuclease will be copied as well; if a gene is present we refer to it as ‘cargo’, as it is ‘driven’ by the endonuclease through the population.

**Fig. 1.**
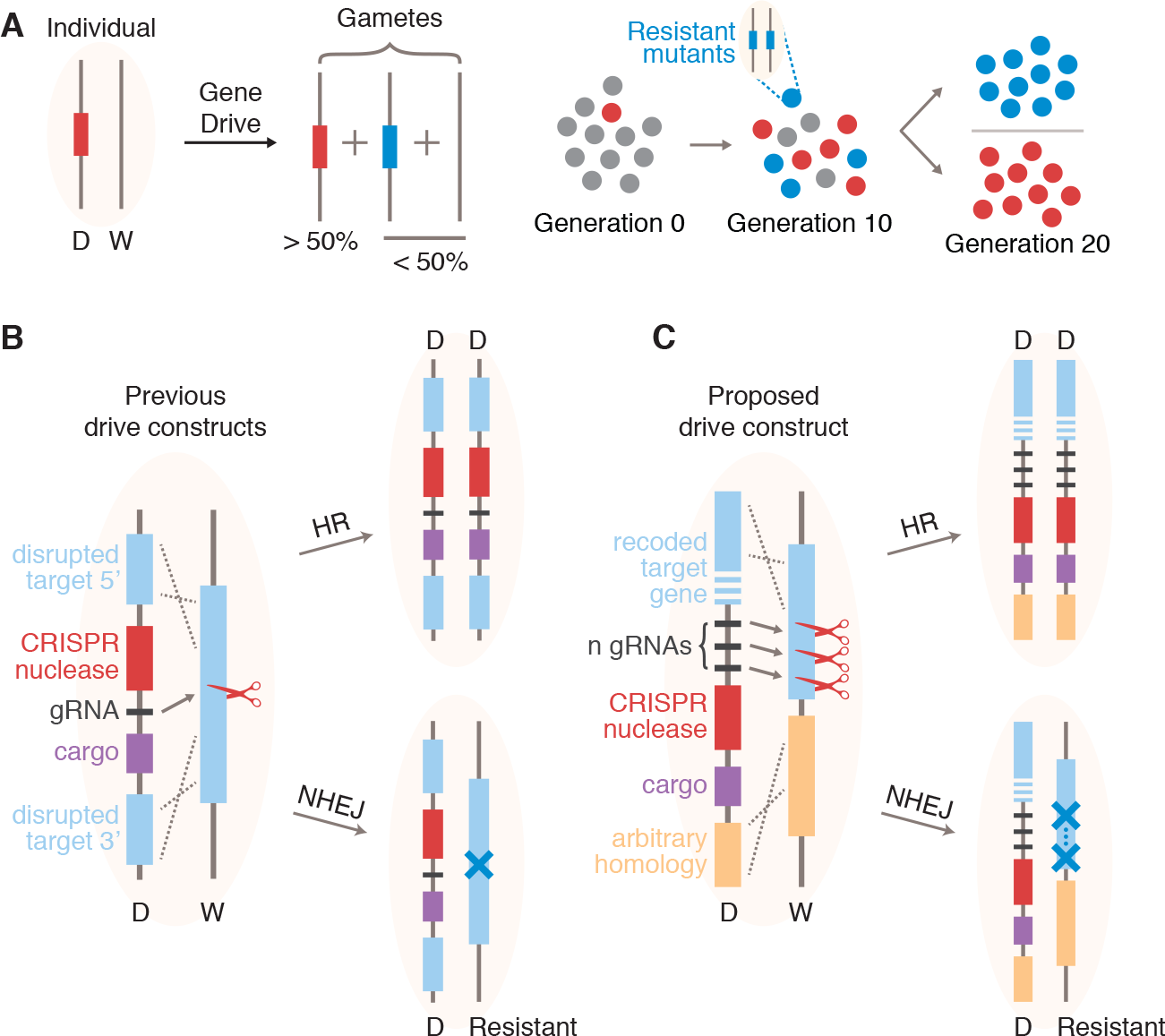
CRISPR gene drive inheritance and spread in wild populations. (**A**), Inheritance and spread of a gene drive construct, D, in a population of individuals homozygous for the wild-type, W. In the late germline, the drive construct induces a double-strand break at its own position on the homologous chromosome which is repaired either by homologous recombination (HR), converting the individual to a DD homozygote, or by non-homologous end joining (NHEJ), producing a small insertion / deletion / substitution mutation at the cut site which results in a drive resistant allele. There is also the possibility of no modification, in which case the W allele remains unchanged. This mechanism can lead to rapid spread of the gene drive in a population or the spread of resistant alleles, depending on their relative fitness effects. (**B**), To achieve this mechanism, previously demonstrated drive constructs are inserted at some target sequence (blue) and carry a CRISPR nuclease (e.g., Cas9) with a guide RNA (gRNA), as well as a “cargo gene” which can be chosen arbitrarily for the desired application. Disruption of the target sequence must be nearly neutral for the drive to spread. (**C**) Our proposed construct reconstitutes the target gene after cutting—so an essential gene can be chosen as the target to select against resistant alleles—and employs multiple (*n*) gRNAs.

Though originally proposed over a decade ago (*1*), the chief technical difficulty of this approach—inducing easily programmable cutting at arbitrary target sites—has only recently been overcome by the discovery and development of the CRISPR/Cas9 genome editing system (*23–27*). Briefly, Cas9 is an endonuclease whose target site is prescribed by an independently expressed guide RNA (gRNA) via a 20-nucleotide protospacer sequence. Because virtually any position in a genome can be uniquely targeted by Cas9, so-called RNA-guided gene drive elements can be constructed by simply inserting a suitable sequence encoding both Cas9 and gRNA(s).

Recent studies have demonstrated highly functional CRISPR gene drive elements in mosquitoes (*5, 6*), yeast (*7*), and fruit flies (*8*). In each case, the basic construct consists of a copy of Cas9 with a single corresponding gRNA and cargo sequence (Fig. 1B). Despite drive inheritance of about 95% on average in the published studies (compared to 50% expected by Mendelian inheritance), the evolutionary stability of these constructs in large populations has been debated due to the potential emergence of drive resistance within a population (*1, 2, 21*). A resistant allele is anticipated to arise whenever the cell repairs the drive-induced DSB using non-homologous end joining (NHEJ) instead of HR, a process which typically introduces a small insertion or deletion mutation at the target sequence. Because the reported constructs cut only at a single site, a large fraction of NHEJ events will create drive-resistant alleles which could prevent the construct from spreading to the entire population (Fig. 1B).

Drive resistance was first mathematically studied in the context of single-cutting homing endonuclease-based drive elements (*21*). There, it was concluded that drive is most effective when the fitness cost of the drive is low and the fitness cost of resistance is high (see SM Section 1 for a description of that work). Unfortunately, in the drive constructs reported thus far, these two requirements are fundamentally at odds: the fitness cost of resistance arises from disruption of the target sequence, but the drive copies itself precisely by disrupting the target sequence.

Here we study the evolutionary dynamics of an alternative drive architecture (*2*) which decouples these effects by rescuing function of the target gene, but only if the drive cassette is successfully copied. This is accomplished by targeting multiple sites within the 3’ end of a gene for cutting by the drive and including a completely genetically recoded (*28, 29*) copy of this 3' target sequence in the drive construct (Fig. 1C). The 3′ UTR of the gene is also replaced with an equivalent sequence in order to remove all homology between the cut sites and the drive components, which ensures that the drive cassette is copied as a single unit. If repair occurs by HR, the target gene is restored to functionality as the drive is copied. But if repair occurs by NHEJ, then the target gene is mutated, potentially resulting in a knockout and a corresponding loss of fitness. Using this design, drive resistance can be selected against by simply choosing an essential or even haploinsufficient gene as the drive target. In addition, the construct employs multiple gRNAs. The use of multiple gRNAs offers two important benefits with respect to resistance: (i) all gRNA target sites must be mutated or lost before a single allele becomes drive-resistant, and (ii) if cutting occurs at two or more gRNA target sites simultaneously, then the intervening DNA sequence is lost, resulting in a large deletion and a knockout of the target gene. This is in contrast to single-cutting constructs, where a knockout can be avoided by an in-frame indel or substitution mutation.

To study this construct, we formulated a deterministic model (SM Sections 2, 7) which considers the evolution of a large population of diploid organisms and focuses on a specific locus with 2*n*+2 alleles (Fig. 2A). First, there are the wild-type (W) allele and the gene drive allele with *n* gRNAs (D). There are then *n* distinct ‘cost-free resistant’ alleles which are resistant to drive-induced cutting at 1, 2, …, *n* target sites but are otherwise identical to the wild-type (denoted S_1_,S_2_,…,S_*n*_). These could arise via, for example, mutations which block cutting by disrupting the gRNA target sequences but do not cause a shift in the reading frame. Finally, there are *n* distinct ‘costly’ resistant alleles which have fitness effects which are distinct from those of the wild-type (denoted R_1_,R_2_, …,R_*n*_). Only the alleles S_*n*_ and R_*n*_ are fully resistant to cutting by the drive. We also refer to the wild-type allele as S_0_ for notational convenience. Finally, we say that individuals having genotype AB, where A and B are any of the alleles above, have fitness *f*_AB_ (alternatively, genotype AB is associated with a *cost* 1-*f*_AB_) and produce gametes having haplotype C with probability *p*_AB,C_. Note that these probabilities *p*_AB,C_ abstract all individual-level drive dynamics and are agnostic to the mechanism which produces drive. We allow these parameters to be arbitrary for our analytical calculations and derive corresponding results which hold for any underlying drive mechanism—including both the previous drive constructs and the new ones considered here.

**Fig. 2.**
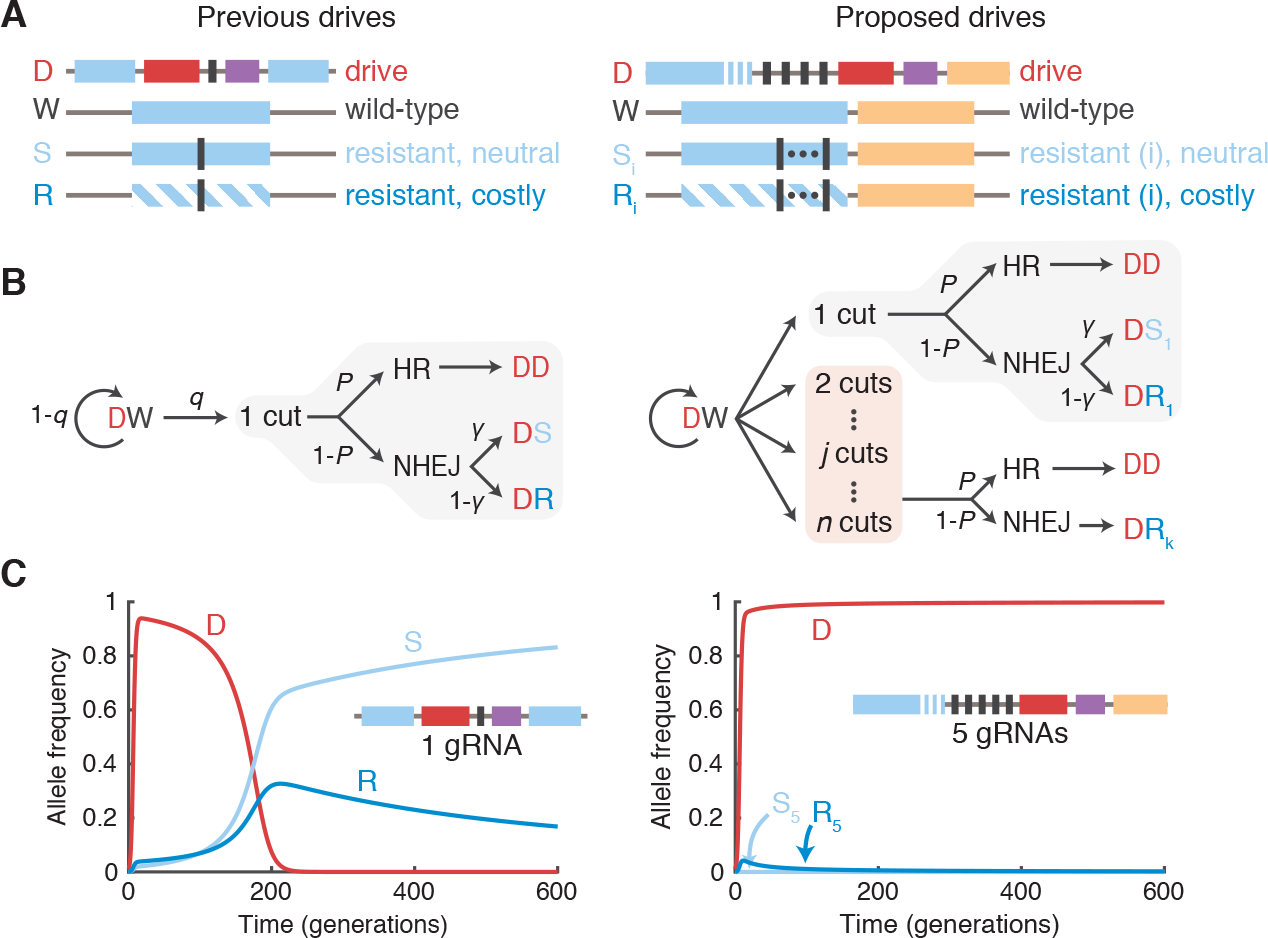
Modeling framework and representative simulations. (**A**) We consider 2*n*+2 alleles, where *n* is the number of drive target sites (prescribed by CRISPR gRNAs): the drive construct (D), the wild-type (W), *n* ‘neutral’ resistant alleles (S_*i*_), and *n* ‘costly’ resistant alleles (R_*i*_). Previous drives (left) have employed one target site, while our proposed drives employ multiple (right). (**B**) Conversion dynamics within DW germline cells during early gametogenesis. Cutting occurs at each susceptible target independently with probability *q*. Then repair occurs by HR with probability *P* or by NHEJ with probability 1-*P*. In the case of a single cut (light grey), repair produces a functional target gene with probability y or a non-functional target with probability 1-γ. Two or more cuts (light red) certainly produce non-functional targets after NHEJ repair. (C) Representative simulations using high cutting and HR probabilities (*q* = *P* = 0.95), for an initial drive release of 1% in a wild-type population, with γ = 1/3. Fitness parameters are (left) *f*_SS_ = *f*_SR_ = 1; *f*_SD_ = 95%; *f*_RR_ = 99%; *f*_DD_ = *f*_DR_ = (99% × 95%) = 94.1%, where S refers to neutral alleles (either S or W), and (right) *f*_SS_ = *f*_SR_ = 1; *f*_SD_ = *f*_DD_ = *f*_DR_ = 95%; *f*_RR_ = 1%, where S and R refer to alleles W, S_1_, …, S_5_ and R_1_, …, R_5_, respectively.

For numerical simulations, we further consider a mechanistic model which explicitly describes the mechanism of drive in individuals (Fig. 2B, Supplementary Material Section 7.3). We assume that, in the germline of an individual which is heterozygous for a drive construct and a susceptible allele (DS_*i*_, where 0 <= *i* < *n* or DR_*i*_, where 1 <= *i* < *n*), each susceptible target site undergoes cutting independently with probability *q*. If there is at least one cut, then HR occurs with probability *P*, while NHEJ occurs with probability 1-*P*. If HR occurs, then the cell is converted to a drive homozygote. But if NHEJ occurs, there are a few possibilities, depending on the number of cuts.

If there is exactly one cut, then one gRNA target is lost on the susceptible allele. If the susceptible allele was initially functional (S_*i*_), then with probability γ it retains function and converts to S_*i*+1_, otherwise it loses function and converts to R_*i*+1_. We assume that the parameter γ is the probability that the reading frame is unaffected, so γ = 1/3. If the susceptible allele is initially nonfunctional (R_*i*_) then we assume that it cannot regain function, so it converts to R_*i*+1_.

If there are two or more cuts, then all *j* susceptible gRNA targets between and including the outermost targets in the locus are lost (2 <= *j* <= *n-i*). The resulting allele is certainly nonfunctional and thus converts to R_i_+j. For simplicity, we assume that the *i* resistant targets are uniformly distributed among the *n* total sites in order to determine a probability distribution for the number of targets lost. We assume that sequential cutting and repair events do not occur.

Now we address two fundamental questions: whether a CRISPR gene drive will invade a resident wild-type population and, if so, whether it will be evolutionarily stable (*30*). We begin with the former. We find that a CRISPR gene drive will invade a wild population if: 
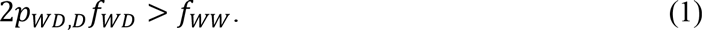

A derivation of this result can be found in Supplementary Material (Sections 3, 7.1). For the drive to spread when initially rare, the advantage from inheritance biasing (*p*_WD,D_)—typically about 95% in published studies—must overcome the difference in fitness between the drive/wild-type heterozygote (*f*_WD_) and the wild-type (*f*_ww_). Note that this condition holds in the context of drive resistance and is agnostic to individual-level drive dynamics and thus applies both to previous drive architectures and our proposed architecture. Indeed, Eq. 1 explains the apparent success of CRISPR drive constructs reported in the literature (*5–8*), which easily invade wild-type laboratory populations—or would be predicted to do so after optimization of drive expression: over short timescales, drive resistance is rare and thus does not affect the dynamics.

However, over longer time scales, NHEJ-mediated resistance will dramatically affect the dynamics. we find that a resident drive population is stable against invasion by resistant alleles only if: 
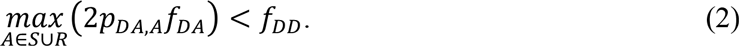

Here the maximization is over all non-drive alleles S_0_, …, S_*n*_, R_*i*_, …, R_*n*_. Intuitively, the drive is stable if and only if no other allele can invade, and each drive allele has an invasion condition identical in form to Eq. 1 (SM Sections 4, 7.2).

Disconcertingly, Eq. 2 suggests that drive constructs are necessarily unstable in sufficiently large populations. An individual which is heterozygous for the drive and the fully-resistant cost free allele S_*n*_ has probability *p*_DSn,Sn_ = ½ of producing an S_*n*_ gamete, and this individual has fitness equivalent to (or potentially greater than) the drive/wild-type heterozygote. Thus if the drive construct has lower fitness than the wild-type, and if the fully-resistant cost-free allele has a nonzero rate of production in the population, the latter will certainly invade a resident drive population. This is especially problematic for highly deleterious population suppression drives, as in (*6*), which have low fitness relative to the wild-type and cost-free resistant alleles.

Population alteration drives (sometimes referred to as replacement drives) might not require long-term persistence in a population to produce their desired effect. Indeed, some applications might still be successful as long as the drive construct attains and persists at a sufficiently high frequency in the population over some length of time.

To quantify the relative effectiveness of the two drive architectures, we considered three quantities: (i) the maximum frequency achieved by a drive construct released in a wild population, (ii) the time required for a drive construct to attain 90% of its maximum frequency, and (iii) the frequency of the drive construct after 200 generations, roughly the longest relevant timescale for a typical application. We computed these quantities numerically for drives featuring cutting and HR probabilities consistent with average drive inheritance rates observed in previous fruit fly (*8*) and mosquito (*5, 6*) experiments (*q* = *P* = 0.95, corresponding to a drive inheritance rate of 95.1% from DW individuals).

Our results suggest that, as anticipated from Eq. 1, both the previous and proposed drive constructs should spread similarly in the short term immediately following release (Fig. 3A, B, and D). However, over longer timescales, the two constructs undergo dramatically different dynamics. The proposed drive constructs, released at an initial frequency of 1% in a wild population, employing five gRNAs and targeting an essential gene, can attain >99% frequency in a population (Fig. 3B, C) in 10-20 generations (Fig. 3B, D) and remain above 99% for at least 200 generations (Fig. 3B, E). Furthermore, this is seen over a large range of drive fitness costs, up to approximately 30% (Fig. 3C–E). The previously demonstrated constructs, in contrast, attain maximum frequencies between 90% and 95% over a narrower range of fitness values (Fig. 3A, C) and demonstrate significantly reduced stability (Fig. 3E). In particular, previous constructs exceeding 8% fitness cost invariably fall below their initial release frequency in fewer than 200 generations.

**Fig. 3.**
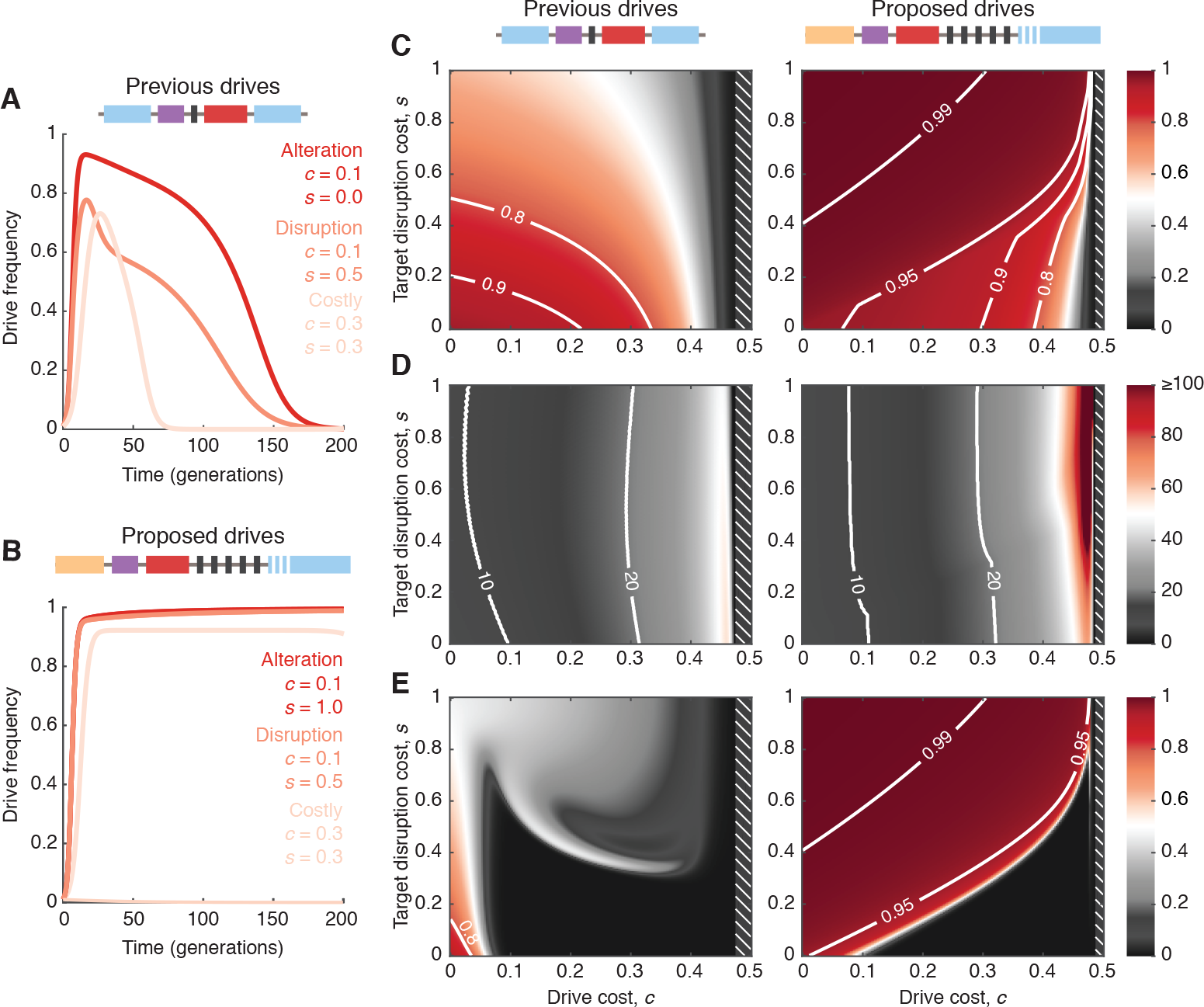
Quantitative comparison of proposed and previously demonstrated drive constructs. (**A** and **B**) Drive frequency over time for three particular scenarios, a low-cost alteration drive carrying a cargo gene (red), a low-cost drive whose aim is to disrupt an important target gene (orange), and a high-cost drive (tan). (**C**) The maximum drive allele frequency (heat) observed in simulations across 200 generations, following an initial release of drive-homozygous organisms comprising 1% of the total population. In white hatched regions, Eq. 1 is not satisfied, so no invasion occurs. (**D**) Generations to 90% of the maximum frequency. (**E**) The frequency of the drive constructs after 200 generations, a measure of stability in the population. Parameters used are: (throughout) *q* = *P* = 0.95. (proposed drives) *n* = 5; *f*_SS_ = *f*_SR_ = 1; *f*_sD_ = *f*_DD_ = *f*_RR_ = 1−*c*;*f*_RR_ = 1−*s*; (previous drives) *n* = 1;*f*_ss_ = *f*_sR_ = 1;*f*_sD_ = 1−*c*; *f*_DD_ = *f*_DR_ = (1−*c*)(1−*s*);*f*_RR_ = 1−*s*, where S and R refer to any alleles S_0_, …, S_*n*_ and R_1_, …, R_*n*_, respectively.

Here we have mathematically shown that previously demonstrated CRISPR gene drives constructed as proofs-of-principle should effectively invade wild populations—consistent with experimental observations—but could have limited utility due to their inherent instability, brought about by their production of resistant alleles. We studied an alternative drive architecture which contains (i) multiple CRISPR guide RNAs which target the 3’ end of a gene, and (ii) a recoded copy of the target gene which is functional but resistant to cutting. We concluded that this architecture substantially improves the stability of CRISPR gene drives.

Another alternative strategy which we have not modeled here would involve multiple independent single-guide drive constructs targeting the same locus. This is conceptually symmetric to the strategy considered here: rather than a single drive with multiple (*n*) gRNAs (“multiple guides”), one might consider multiple (*n*) drives with one gRNA each (“multiple drives”). In this strategy, each independent drive would behave similarly to the previously demonstrated constructs studied here. This strategy would likely outperform the previous strategy, but we anticipate that it would not outperform the multiple guide strategy. This is because, in the multiple drive strategy, each gRNA target can undergo NHEJ-mediated mutation independently, providing stepping-stones to fully-resistant alleles. Furthermore, the multiple drive strategy lacks the benefit of large NHEJ knockouts from multiple simultaneous cuts which help combat cost-free resistance (Fig. 2B, red box), although it would be capable of editing regions unimportant to fitness.

In conclusion, we suggest three concrete design principles for future CRISPR gene drive systems. Constructs will maximize efficacy and stability if (i) multiple guide RNAs with minimal off-target effects are employed, (ii) disruption of the target locus is highly deleterious, and (iii) any cargo genes are as close to neutral as possible.

## Supplementary Material

In this Supplementary Material, we mathematically study the evolutionary dynamics of a CRISPR gene drive construct with *n* guide RNAs. In Section 1, we discuss relevant prior work on homing endonuclease gene drives. In Section 2, we propose a simple model of population genetics of RNA-guided gene drives with multiple guide RNAs, and we analyze the selection pressure acting on an engineered drive construct. In Section 3, we derive a condition for an engineered drive allele to invade a natural population. In Section 4, we derive a condition for a population in which the drive has fixed to resist invasion by either wild-type or drive-resistant alleles. In Section 5, we derive equations for interior equilibria permitted by our system. In Section 6, we present numerical examples of the system's dynamics. In Section 7, we extend the model from Section 2 to include the effects of “neutral resistance”.

### 1 Previous work on homing endonuclease gene drives

Deredec et al. (2008) (Ref. (21) in the main text) mathematically investigates the evolutionary dynamics of homing endonuclease gene drives. The authors begin with a two-allele model precluding resistance, consisting of a wild-type allele and a gene drive allele (pp. 2014-2016 of Deredec et al. (2008)). The model implicitly considers a single guide RNA because it was motivated by earlier single-target homing endonuclease genes. In their notation, *p* is the frequency of the wild-type allele, and *q* is the frequency of the drive allele. The authors assume Hardy-Weinberg proportions at all times, and they write a recurrence for *q*: 
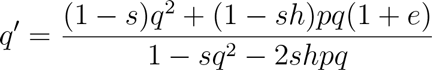

Here, *s* is the fitness cost associated with a drive homozygote, *sh* is the fitness cost associated with a drive/wild-type heterozygote, and *e* is the probability that the HEG copies itself onto the homologous chromosome (“homes”).

The authors identify that there are three possible fixed points: 
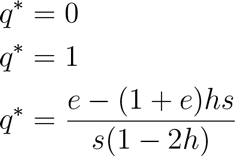

To obtain the invasion condition for the drive allele, the authors solved 
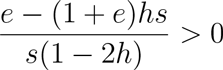

They obtain 
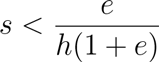

Intuitively, the fitness cost, *sh*, of a drive/wild-type heterozygote must be less than a monotonically increasing function of the homing rate, *e*, for the homing endonuclease gene to spread when rare. Low fitness costs of the drive and high homing rates facilitate the invasion of the drive. More specifically, the authors show that, if the drive/wild-type heterozygote has fitness close to the wild-type (i.e., *h* close to zero), then the drive invades and fixes (if *s* is small relative to *e*), coexists with the wild-type allele (if *s* is comparable in magnitude to *e*), or does not invade and is unstable (if *s* is large relative to *e*). The authors also show that, if the drive/wild-type heterozygote has fitness close to the drive homozygote (i.e., *h* close to one), then the drive invades and fixes (if *s* is small relative to *e*), is bistable with the wild-type allele (if *s* is comparable in magnitude to *e*), or does not invade and is unstable (if *s* is large relative to *e*). These are important insights into the evolutionary dynamics of homing endonuclease gene drives.

Deredec et al. then extend their model to consider also a single resistant allele (pp. 2018-2019 of Deredec et al. (2008)). In their notation, *p* is the frequency of the wild-type allele, *q_H_* is the frequency of the drive allele, and *q_M_* is the frequency of the misrepaired (resistant) allele. The authors assume Hardy-Weinberg proportions at all times, and they write recurrences for *q_H_* and *q_M_*: 
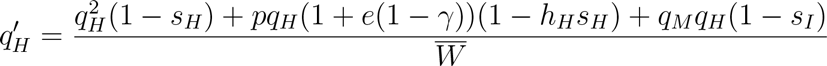

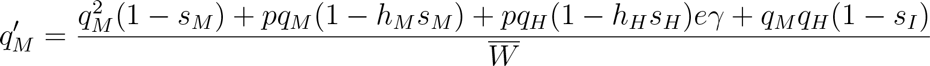

Here, 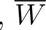 is the mean fitness of the population, and γ is the probability of misrepair.

The authors then consider a variety of special cases and make observations about each. A general theme is that low misrepair rates, high fitness of the drive, and low fitness of resistance alleles all act to improve drive spread. These are crucial points for understanding the evolutionary dynamics of homing endonuclease genes.

For a classic homing endonuclease gene drive, the latter two properties—high fitness of the drive and low fitness of resistance alleles—are naturally difficult to reconcile with each other, as we describe in the main text. Since cost-free resistance to a drive construct arises, alternative drive designs are necessary for effective genome editing. The recently developed CRISPR/Cas9 genome editing technology facilitates targeting arbitrary locations in a genome, greatly expanding the creative potential for manipulating wild populations. While CRISPR/Cas9 constructs offer enhanced opportunities for constructing functional gene drives, they also inevitably exhibit more complex dynamics that must be firmly understood.

### 2 Model for the evolutionary dynamics of a CRISPR gene drive with *n* guide RNAs

Consider a wild population of diploid organisms. Our aim is to manipulate the population by modifying a particular locus which may be important, for example, for the organism's survival, reproduction, disease transmission, etc. Using CRISPR/Cas9 genome editing techniques, one can engineer a CRISPR gene drive with *n* guide RNAs to target this locus. See the main text and corresponding Fig. 1 for specific discussion of our proposed design.

To describe the evolutionary dynamics of such a construct, we consider a drive allele, D, a wild-type allele, 0, and *n* resistance alleles, *i* (with 1 ≥ *i* ≥ *n*). (In the main text, we use the notation “W” for a wild-type allele instead of “0”. The notation “0” is more natural for doing calculations.) There are (*n* + 2) + (*n* + 2)(*n* + 1)/2 possible individual genotypes in the population: *ij* (with 0 ≥ *i ≥ n* and 0 ≥ *j* ≥ *n*), *iD* (with 0 ≥ *i* ≥ *n*), and *DD*. The drive mechanism works as follows:

Consider a type 0*D* individual; one allele is wild-type, and the other allele is the drive. There are *n* guide RNAs and therefore *n* targets for the drive to cut. At meiosis, the drive can cut any number of targets between 0 and *n*. If the drive cuts no targets, then the individual remains with genotype 0*D*. If the drive cuts *k* targets (with 1 ≥ *k* ≥ *n*), then one of several things can happen: One possibility is that homologous recombination copies the drive allele onto the damaged chromosome, so that the individual's genotype becomes *DD*. This is how the drive construct effects its spread through a population. Another possibility is that non-homologous end joining repairs the damaged chromosome without restoring the lost targets, so that the individual's genotype becomes *iD* (with 1 ≥ *i* ≥ *k*). This is how resistance to the drive construct emerges. Yet another possibility is that non-homologous end joining perfectly repairs the damaged chromosome, so that the individual's genotype remains 0*D*.

The drive allele can effect its spread as long as there is at least one remaining target. For example, in an individual with genotype *iD*, the drive can cut at any number, *k*, of the *n* − *i* remaining targets (so that 1 ≥ *k* ≥ *n* − *i*). After cutting, the individual can become homozygous in the drive allele (*DD*), the individual can lose additional targets by acquiring genotype *jD* (with *i* + 1 ≥ *j* ≥ *i* + *k*), or the individual can remain with genotype *iD*.

Using these rules, we can formally express the rates at which each of the *n* + 2 types of gametes are produced in terms of the frequencies of individuals in the population. We denote by *F_D_* (*t*) the rate (at time *t*) at which drive gametes (*D*) are produced by individuals in the population. We denote by *F_i_* (*t*) the rate (at time *t*) at which wild-type gametes (*i* = 0) or gametes with varying levels of resistance (1 ≥ *i* ≥ *n*) are produced by individuals in the population. We have 
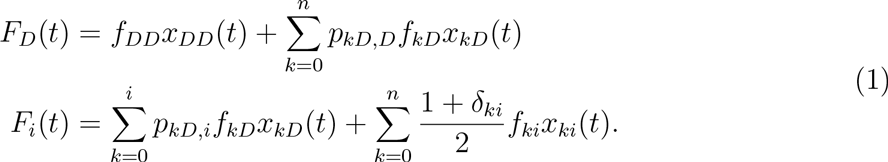
*δ_ki_* is the Kronecker delta. We use the following notation: *x_ki_*(*t*) denotes the frequency of individuals (at time *t*) with only wild-type or resistance alleles, *x_kD_*(*t*) denotes the frequency of individuals (at time *t*) with one wild-type or resistance allele and one drive allele, and *x_DD_* (*t*) denotes the frequency of individuals (at time *t*) that are homozygous in the drive allele. (We define *x_ki_*(*t*) for *k* ≠ *i* and *x_kD_*(*t*) such that the ordering of the indices does not matter, i.e., *x_ki_*(*t*) = *x_ik_*(*t*) is the total frequency of individuals with either genotype *ki* or genotype *ik*, and *x_kD_*(*t*) = *x_Dk_*(*t*) is the total frequency of individuals with either genotype *kD* or genotype *Dk*.) *f_ki_* denotes the fitness of individuals with only wild-type or resistance alleles, *f_kD_* denotes the fitness of individuals with one wild-type or resistance allele and one drive allele, and *f_DD_* denotes the fitness of individuals that are homozygous in the drive allele. *p_kD,D_* denotes the probability that an individual of genotype *kD* produces a *D* gamete. *p_kD,i_* denotes the probability that an individual of genotype *kD* produces an *i* gamete. From conservation of probability, we have the following identity: 
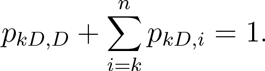

Notice that a type *nD* individual is fully resistant to being manipulated by the drive construct; such a fully resistant individual shows standard Mendelian segregation in its production of gametes. Thus, we have 
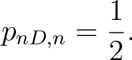

We understand Equations (1) as follows: Type *DD* individuals only produce type *D* gametes, hence the term *f_DD_x_DD_*(*t*) in the equation for *F_D_*(*t*). Type *kD* individuals produce type *D* gametes with probability *p_kD,D_*, hence the terms *p_kD,D_f_kD_x_kD_*(*t*) in the equation for *F_D_* (*t*). Type *kD* individuals produce type *i* gametes with probability *p_kD,i_*, hence the terms *p_kD,i_f_kD_x_kD_*(*t*) in the equation for *F_i_*(*t*). Type *ki* individuals produce type *i* gametes with probability 1 if *k* = *i* or with probability 1/2 if *k* ≠ *i* hence the terms [(1 + *δ_ki_*)/2]*f_ki_x_ki_*(*t*) in the equation for *F_i_*(*t*).

The selection dynamics are modeled by the following system of equations: 
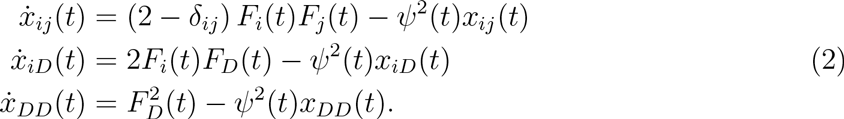

Here, an overdot denotes the time derivative, *d/dt*. In formulating the population dynamics, we assume random mating; i.e., two random gametes meet to form a new individual. Notice that the products (2 − *δ_ij_*)*F_i_*(*t*)*F_j_*(*t*), 2*F_i_*(*t*)*F_D_*(*t*), and 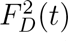 in Equations (2) represent the pairings of the different types of gametes to make new offspring. The quantity *ψ^2^*(*t*) represents a density-dependent death rate for the individuals in the population.

At any given time, *t*, we require that the total number of individuals sums to one: 
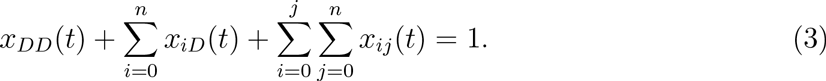

To enforce this density constraint, we set 
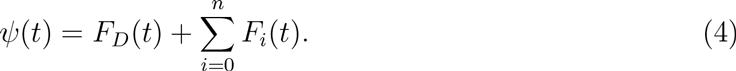

Throughout this SM, we choose to work in the framework of continuous time (Equations (2)), since we feel that this approach simplifies the mathematical analysis. In much of the remainder of this SM, we omit explicitly writing the time dependence on dynamical quantities for notational convenience.

### 3 Invasion of the drive construct

Consider a wild-type population in which all individuals have genotype 00. We perturb the wild-type population by introducing a small amount of the drive allele, *D*. What happens? Does the drive allele catalyze its own spread in the population, or is it eliminated?

For a perturbation to a wild-type population, we write the frequencies of the individual genotypes as 
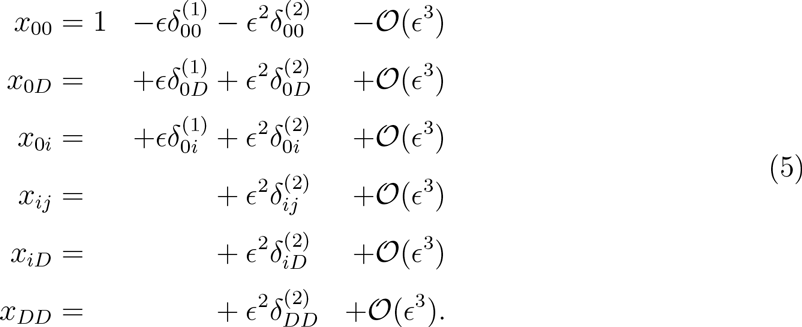

In Equations (5), it is implied that 1 ≥ *i* ≥ *n*. The expansions (5) are understood as follows. The frequency of the wild-type allele is approximately one, since we only introduce a small amount of the drive allele. The frequency of the drive allele is of order *ϵ* ⪡ 1. The small number of 0*D* individuals in the population also produce resistance alleles, and the frequency of these resistance alleles shortly after the perturbation is also small (i.e., of order *ϵ* ⪡ 1). Notice that:

- New type 00 individuals are produced by pairing two wild-type gametes (each at frequency 𝒪(1)), so new type 00 individuals are generated at a rate of order 1.
- New type 0*D* individuals are produced by pairing a wild-type gamete (at frequency 𝒪(1)) and a drive gamete (at frequency 𝒪(*ϵ*)), so new type 0*D* individuals are generated at a rate of order *ϵ*.
- New type 0*i* individuals (for 1 ≤ *i* ≤ *n*) are produced by pairing a wild-type gamete (at frequency 𝒪(1)) and a resistant gamete (at frequency 𝒪(*ϵ*)), so new type 0*i* individuals are generated at a rate of order *ϵ*.
- New type *ij* individuals (for 1 ≤ *i* ≤ *n* and 1 ≤ *j* ≤ *n*) are produced by pairing two resistant gametes (each at frequency 𝒪(*ϵ*)), so new type *ij* individuals are generated at a rate of order *ϵ*^2^.
- New type *iD* individuals are produced by pairing a resistant gamete (at frequency 𝒪(*ϵ*)) and a drive gamete (at frequency 𝒪(*ϵ*)), so new type *iD* individuals are generated at a rate of order *ϵ*^2^.
- New type *DD* individuals are produced by pairing two drive gametes (each at frequency 𝒪(*ϵ*)), so new type *DD* individuals are generated at a rate of order *ϵ*^2^.

Also, notice that a nonzero amount of the drive allele and the resistant allele are each produced at order *ϵ*^2^ by type *ij, iD*, and *DD* individuals, so there also exist terms of order *ϵ*^2^ in the expansions for *x*_0*D*_ and *x*_0*i*_. Hence, we arrive at the expansions (5).

Note that (5) and (3) impose a constraint on the 𝒪(*ϵ*) terms in the genotype frequencies: 
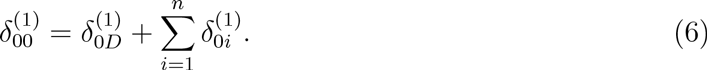

Also, note that (5) and (3) impose a constraint on the 𝒪(*ϵ*^2^) terms in the genotype frequencies: 
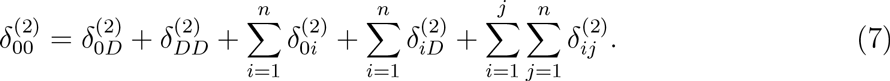

Substituting (4), (1), (5), and (6) into the equation for *x*_0*D*_ in (2), we obtain 
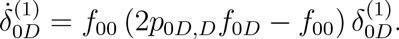

The drive allele invades a wild-type population if 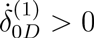, i.e., if 
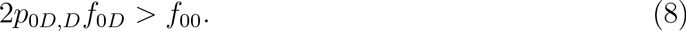

### 4 Stability of the drive construct

Consider a population in which the drive construct has fixed, so that all individuals have genotype *DD*. We perturb the *DD* population by introducing a small amount of the wild-type allele, 0. What happens? Is the *DD* population stable to perturbations, or does the wild-type allele or one of the resistance alleles invade the population?

For a perturbation to a population in which the drive construct has fixed, we write the frequencies of the individual genotypes as 
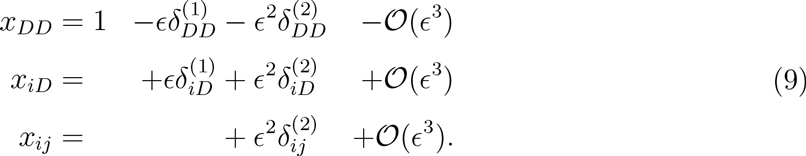

The expansions (9) are understood as follows. The frequency of the drive allele is approximately one, since we only introduce a small amount of the wild-type allele. The frequency of the wild-type allele is of order *ϵ* ⪡ 1. The small number of 0*D* individuals in the population also produce resistance alleles, and the frequency of these resistance alleles shortly after the perturbation is also small (i.e., of order *ϵ* ⪡ 1). Notice that:

- New type *DD* individuals are produced by pairing two drive gametes (each at frequency 𝒪(1)), so new type *DD* individuals are generated at a rate of order 1.
- New type *iD* individuals (for 0 ≤ *i* ≤ *n*) are produced by pairing a non-drive gamete (at frequency 𝒪(*ϵ*)) and a drive gamete (at frequency 𝒪(1)), so new type *iD* individuals are generated at a rate of order *ϵ*.
- New type *ij* individuals (for 0 ≤ *i* ≤ *n* and 0 ≤ *j* ≤ *n*) are produced by pairing two non-drive gametes (each at frequency 𝒪(*ϵ*)), so new type *ij* individuals are generated at a rate of order *ϵ*^2^.

Also, notice that a nonzero amount of the non-drive alleles are produced at order *ϵ*^2^ by type *ij* individuals, so there also exist terms of order *ϵ*^2^ in the expansions for *x_iD_*. Hence, we arrive at the expansions (9).

Note that (9) and (3) impose a constraint on the 𝒪(*ϵ*) terms in the genotype frequencies: 
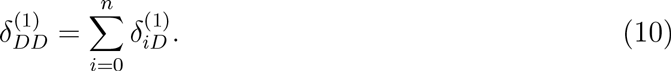

Also, note that (9) and (3) impose a constraint on the 𝒪(*ϵ*^2^) terms in the genotype frequencies: 
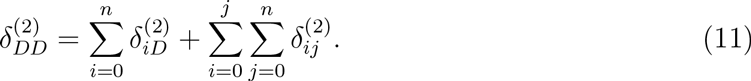

Substituting (4), (1), (9), and (10) into the equations for *X*in (2), we obtain 
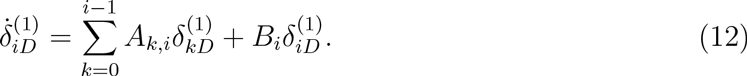

Here, we use the shorthand notation 
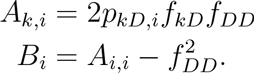

The solution to (12) is 
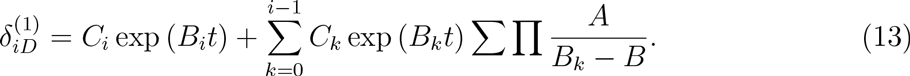

The quantities *C_i_* in (13) are arbitrary constants of integration. The sum of products in (13) is equal to 
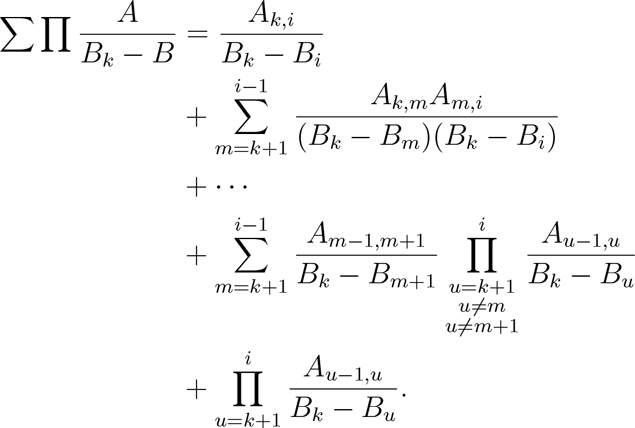

Notice that the time dependence in (13) appears only in the exponential factors. If all *B_i_* < 0, then all 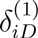 approach zero in the long-time limit, and, from (10), we have that 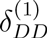 approaches zero in the long-time limit. Therefore, if *B_i_* < 0 for all values of 0 ≥ *i* ≥ *n*, then the drive construct is evolutionarily stable.

If, instead, *B_i_* > 0 for at least one value of *i*, then 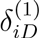 has a term whose magnitude grows exponentially in time. The leading-order (in *ϵ*) terms in the expansions for *x_iD_* in (9) are necessarily positive. Therefore, if the condition *Bi* > 0 is satisfied for at least one value of *i*, then 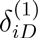 is positive and grows exponentially in time; i.e., the *DD* population is unstable to perturbations.

The resulting condition is that the *DD* population is stable to perturbations with a wild-type allele if 
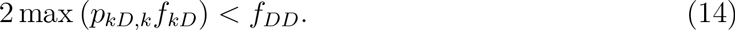

#### 4.1 Completely recessive fitness cost for a resistance mutation

Here, we consider a special case in which the fitness cost associated with having resistance to the drive is completely recessive. If the fitness of each heterozygote with a resistance allele, *f_kD_*, exactly equals *f_DD_* for all *k*, then is the *DD* population stable to perturbations? We expect that *p_kD,k_* < 1/2 for all 0 ≤ *k* < *n*. Therefore, if *f_kD_* = *f_DD_* for all *k*, then the inequality (14) is satisfied for all *k* < *n* and becomes an equality for *k* = *n*.

All resistance alleles with at least one target (0 ≤ *k* < *n*) are removed from the population by selective forces. We must focus on the fully resistant allele, *n*. To probe the stability of the *DD* population, we substitute (4), (1), (9), (10), and (11) into (2), and we keep terms that are 𝒪(*ϵ*^2^). We have 
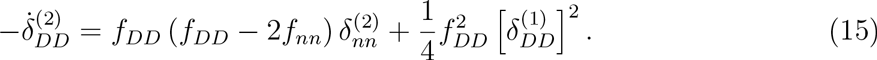

We also have 
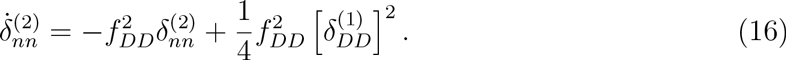

We can integrate (16). We get 
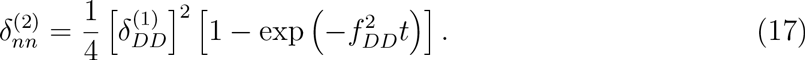

We are interested in the regime 1 ⪡ *t* ⪡ *ϵ*^−1^. We must consider the sign of 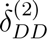 at large times *t* ⪢ 1 but before the terms in (9) become similar in magnitude. Our condition for stability of the *DD* population is therefore 
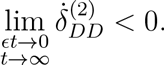

On a short time scale, the exponential in the solution for 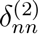 will approach zero. Substituting (17) into (15) and simplifying, we see that the *DD* population is stable to perturbations if 
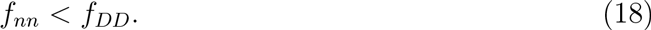

### 5 Interior equilibria

A drive construct increases in frequency when rare if Equation (8) is satisfied. A drive construct that has already fixed is stable to perturbations if Equation (14) is satisfied (or if Equation (18) is satisfied for the case of a completely recessive fitness cost for resistance). But if a small amount of the drive construct is introduced into a wild-type population, then does the drive spread completely to fixation?

To answer this question, it is helpful to know if the model for the drive dynamics, Equations (2), admits an interior equilibrium. Notice that, if all time derivatives are zero, then Equations (2) simplify to 
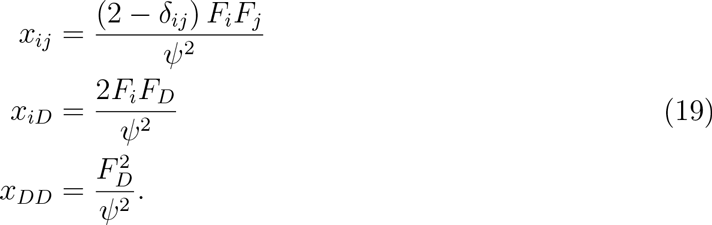

Next, we define *x_i_* to equal the frequency of allele *i* in the population. Thus, *x*_0_ is the frequency of the wild-type allele, and *x_i_* for 1 ≤ *i* ≤ *n* is the frequency of a resistance allele with *i* damaged targets. Also, *x_D_* is the frequency of the drive allele. These allele frequencies can be calculated from the frequencies of individuals of the various genotypes: 
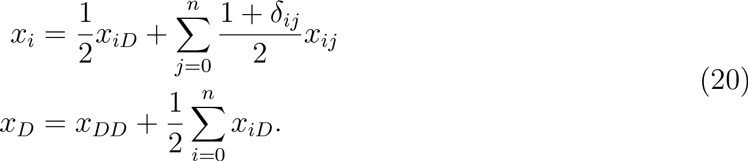

Similarly to Equation (3), the sum of all allele frequencies equals 1 at all times: 
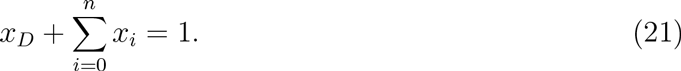

We directly compute the following results: 
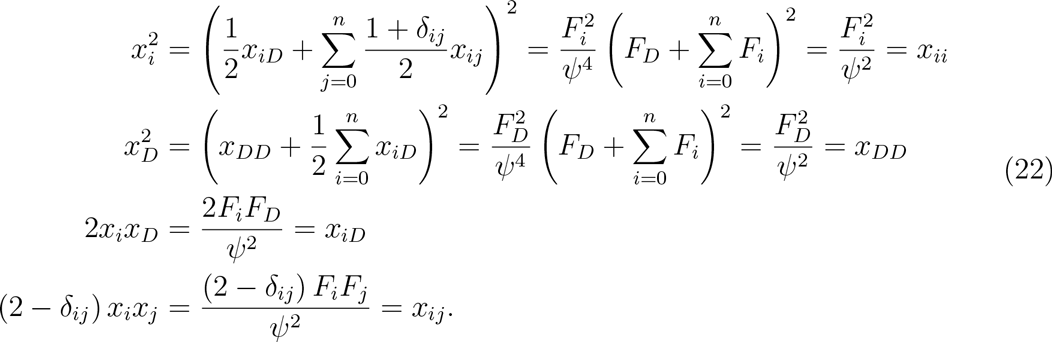

In summary, we obtain 
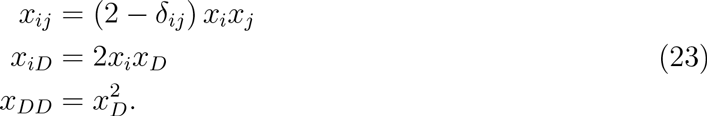

Thus, notice that, at an equilibrium point of the dynamics, the individual frequencies are exactly at Hardy-Weinberg proportions.

From (22), we have that 
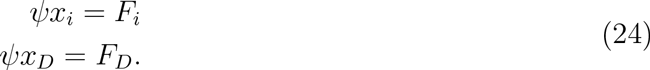

By substituting Equations (1) for *F_i_* and *F_D_* into (24), and substituting (23), we obtain 
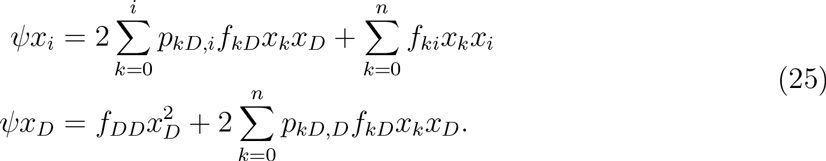

If the drive construct is at a nonzero frequency (*x_D_* > 0), then we can cancel an overall factor of *x_D_* from the second equation of (25): 
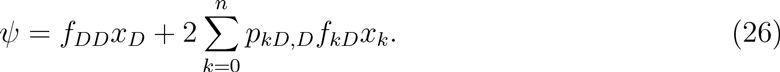

Substituting (26) into the first equation of (25), we have 
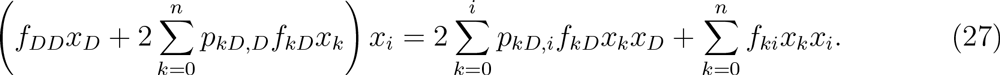

We can also substitute the density constraint, (21), into (27). We obtain 
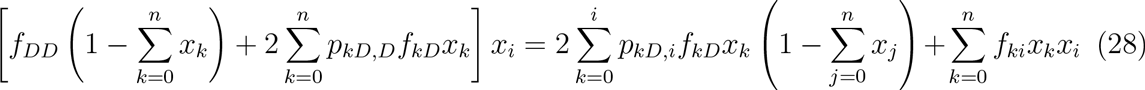

Equations (28) are a set of *n* +1 independent conditions in *n* + 1 variables that must be simultaneously satisfied for each interior fixed point. If Equations (28) cannot be simultaneously solved for a given set of parameter values, then no interior fixed point exists.

#### 5.1 One guide (*n* = 1)

It is instructive to consider the system of equations (28) for the case of a single guide (*n* = 1). Substituting *i* = 0 into (28), we obtain one equation: 
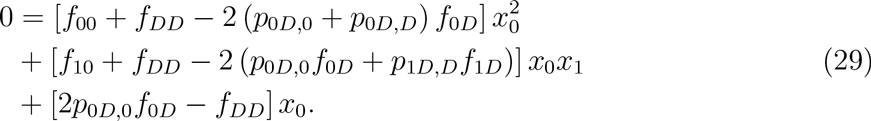

Substituting *i* = 1 into (28), we obtain another equation: 
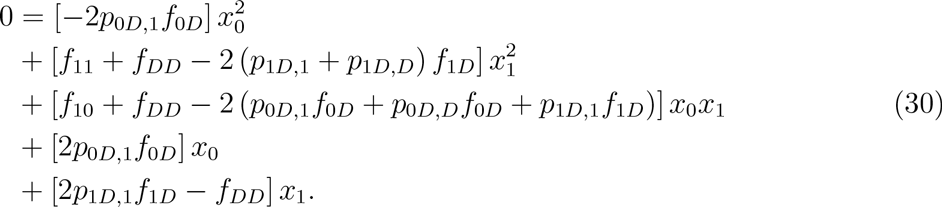

If Equations (29) and (30) cannot be simultaneously satisfied for given values of the parameters *f*_00_, *f_DD_*, *f*_11_, *f*_0*D*_, *f*_1*D*_, *f*_10_, *P*_0*D,D*_, and *p*_0*D*,0_, then no interior fixed point exists.

### 6 Numerical examples

Numerical simulations of Equations (2) are helpful for understanding the evolutionary dynamics of a drive construct. For simplicity, we consider a single guide (*n* = 1), and we choose the following parameter values: 
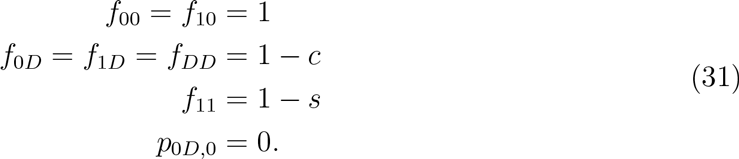

We make the following assumptions: The fitness cost of the drive, *c*, is dominant. The fitness cost of the resistant allele, *s*, is recessive. Also, the drive construct in a 0*D* heterozygote always cuts at the target, and either the drive allele is copied by homologous recombination or resistance emerges. Thus, we have *P*_0*D*,0_, = 0.

In Fig. S1 (a and b), numerical simulations demonstrate evolutionary invasion of the drive construct. For these simulations, the initial condition is *x_AA_* = 1 − 10^−4^ and *x_DD_* = 10^−4^. The relevant condition for determining evolutionary invasion is Equation (8).

- In Fig. S1 (a), we set *p*_0*D,D*_ = 0.75 and *s* = 0.4. From Equation (8), the critical value of *c* for invasion is 1/3. If *c* = 0.34 (green curve), then the drive construct does not invade. If *c* = 0.33 (blue curve), then the drive construct invades.
- In Fig. S1 (b), we set *p*_0*D,D*_ = 0.65 and *s* = 0.3. From Equation (8), the critical value of *c* for invasion is 1/3. If *c* = 0.235 (green curve), then the drive construct does not invade. If *c* = 0.225 (blue curve), then the drive construct invades.

In Fig. S1 (c and d), numerical simulations demonstrate evolutionary stability of the drive construct. For these simulations, the initial condition is *x_DD_* = 1 − 10^−2^ and *x_AA_* = 10^−2^. From (31), notice that the condition (14) becomes an equality. Therefore, the relevant condition for determining evolutionary stability is Equation (18).

- In Fig. S1 (c), we set *p*_0*D,D*_ = 0.75 and *c* = 0.32. From Equation (18), the critical value of *s* for stability is 0.32. If *s* = 0.315 (green curve), then the drive construct is unstable. If *s* = 0.325 (blue curve), then the drive construct is stable.
- In Fig. S1 (d), we set *p*_0*D,D*_ = 0.65 and *c* = 0.2. From Equation (18), the critical value of *s* for stability is 0.2. If *s* = 0.195 (green curve), then the drive construct is unstable. If *s* = 0.205 (blue curve), then the drive construct is stable.

In Fig. S1 (e and f), numerical simulations demonstrate fixation of the drive construct. For these simulations, the initial condition is *x_AA_* = 1 − 10^−4^ and *x_DD_* = 10^−4^. If Equations
(29) and (30) cannot simultaneously be solved numerically, then there is no internal equilibrium.

- In Fig. S1 (e), we set *p*_0*D,D*_ = 0.75 and *c* = 0.32. From numerical analysis of Equations (29) and (30), the critical value of *s* for non-existence of an interior equilibrium is approximately 0.82. If *s* = 0.81 (green curve), then the drive construct reaches an equilibrium frequency that is strictly between 0 and 1. If *s* = 0.82 (blue curve), then the drive construct spreads to fixation.
- In Fig. S1 (f), we set *p_QD_,_D_* = 0.65 and *c* = 0.32. From numerical analysis of Equations (29) and (30), the critical value of *s* for non-existence of an interior equilibrium is approximately 0.29. If *s* = 0.28 (green curve), then the drive construct reaches an equilibrium frequency that is strictly between 0 and 1. If *s* = 0.29 (blue curve), then the drive construct spreads to fixation.

### 7 Neutral resistance

In this section, we present an extension of the model which accounts for the phenomenon of “neutral resistance”. This can occur if non-homologous end-joining results in repair at a cut site which disrupts the recognition sequence of a guide RNA while nonetheless leaving the function of the target gene intact. This can occur, for example, via an in-frame insertion or deletion or synonymous mutation. The resulting allele is similar (with respect to the drive mechanism) to the resistant alleles discussed in previous sections: the repaired target is immune to cutting by its corresponding guide RNA. However, the mutation conferring this resistance is not deleterious.

We represent this scenario by an extension of our original model (Section 2). We consider a drive allele, *D*, *n* “costly” resistant alleles, *R_i_* (with 1 ≤ *i* ≤ *n*), *n* “neutral” resistant alleles, *S_i_* (with 1 ≤ *i* ≤ *n*), and the wild-type allele, *S_0_*. The drive mechanism works as follows (see Figure 2 in the main text for an illustration):

Consider a type *S*_0*D*_ individual; one allele is wild-type, and the other allele is the drive. There are *n* guide RNAs and therefore *n* targets for the drive to cut. At meiosis, the drive can cut any number of targets between 0 and *n*. If the drive cuts no targets, then the individual remains with genotype *S*_0*D*_. If the drive cuts *k* targets (with 1 ≤ *k* ≤ *n*), then one of several things can happen:

- One possibility is that homologous recombination copies the drive allele onto the damaged chromosome, so that the individual's genotype becomes *DD*. This is how the drive construct effects its spread through a population.
- Another possibility is that non-homologous end joining repairs the damaged chromosome. We assume that each of the *k* cut sites are repaired such that they are resistant to later cutting, and the resulting resistant allele is either costly, in which case the individual's genotype becomes *DR_k_*, or cost-free, in which case the individual's genotype becomes *DS_k_*.

The drive allele can effect its spread as long as there is at least one remaining target. In an individual with genotype *R_i_D* or *S_i_D*, the drive can cut at any number, *k*, of the *n* − *i* remaining targets (so that 1 ≤ *k* ≤ *n* − *i*). After cutting, the individual can become homozygous in the drive allele (*DD*), or the individual can lose additional targets by acquiring genotype *R_j_D* or *S_j_D* (with *i* + 1 ≤ *j* ≤ *i* + *k*). We assume that costly resistant alleles *R_i_* cannot convert to cost-free resistant alleles *S_j_*, but cost-free resistant alleles *S_i_* can convert to costly resistant alleles *R_j_*.

**Fig. S1.**
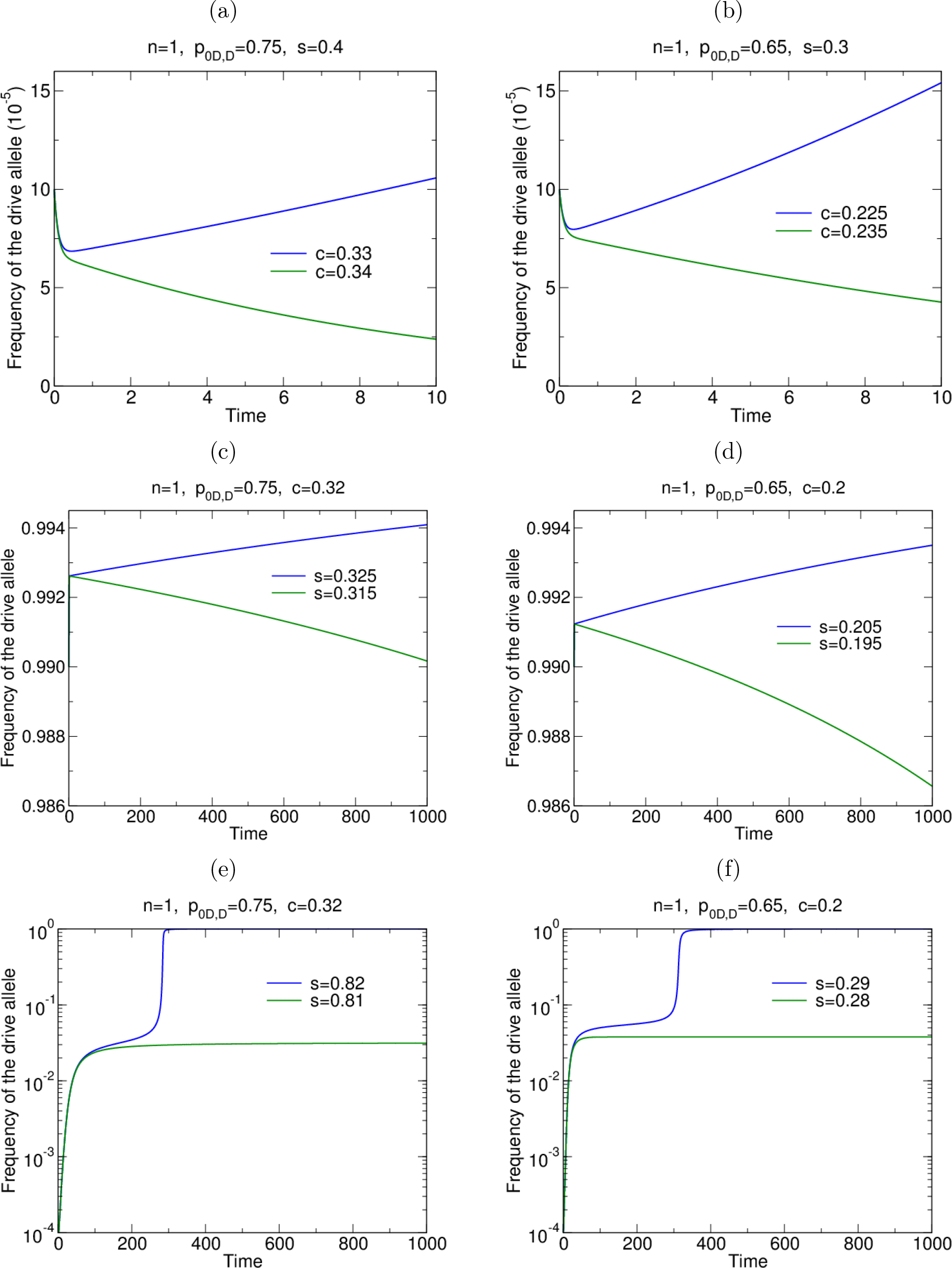
Numerical simulations of the evolutionary dynamics demonstrate the conditions for drive invasion and stability. If no interior equilibrium exists, then an initially rare drive construct spreads to fixation.

Using these rules, we can formally express the rates at which each of the 2*n* + 2 types of gametes are produced in terms of the frequencies of individuals in the population. We denote by *F_D_* (*t*) the rate (at time *t*) at which drive gametes (*D*) are produced by individuals in the population. We denote by *F_R_i__* (*t*) the rate (at time *t*) at which gametes with varying levels of costly resistance (1 ≤ *i* ≤ *n*) are produced by individuals in the population. And we denote by *F_S_i__* (*t*) the rate (at time *t*) at which wild-type gametes (*i* = 0) or gametes with varying levels of cost-free resistance (1 ≤ *i* ≤ *n*) are produced by individuals in the population. We have 
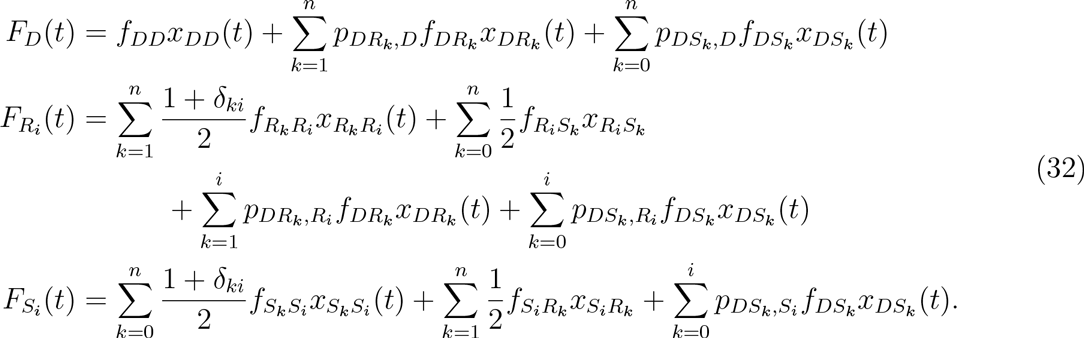

Here, *δ_ki_* is the Kronecker delta. *x_IJ_*(*t*) denotes the frequency of individuals (at time *t*) with genotype *IJ*, where *I,J* = *D, S*_0_, *S*_1_,…, *S_n_, R*_1_,…, *R_n_*. Similarly, *f_IJ_* is the fitness of *IJ* individuals, and *p_IJ,K_* denotes the probability of an individual with genotype *IJ* producing a *K* gamete. From conservation of probability, we have the following identities: 
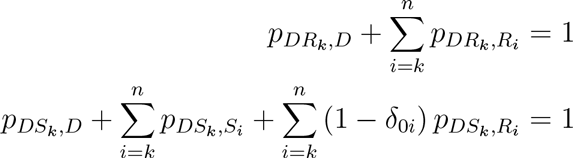

Notice that type *R_n_D* and type *S_n_D* individuals are fully resistant to being manipulated by the drive construct; such a fully resistant individual shows standard Mendelian segregation in its production of gametes. Thus, we have 
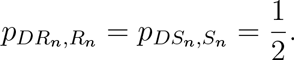

The selection dynamics are modeled by the following system of equations: 
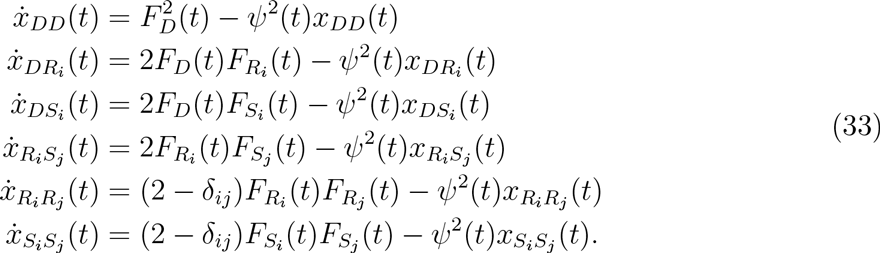

The quantity *ψ*^2^(*t*) represents a density-dependent death rate for the individuals in the population.

At any given time, *t*, we require that the total number of individuals sums to one: 
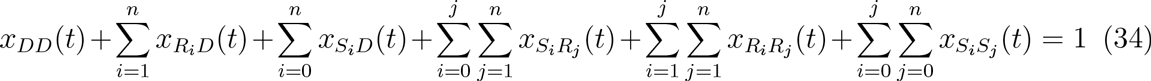

To enforce this density constraint, we set 
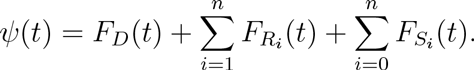

#### 7.1 Invasion of the drive construct

The steps for determining if the drive construct invades when there is neutral resistance are the same as in Section 3. The drive allele invades a wild-type population if 
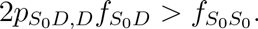

#### 7.2 Stability of the drive construct

The steps for determining if the drive construct is stable when there is neutral resistance are the same as in Section 4. The *DD* population is stable to perturbations with a wild-type allele if 
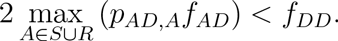

#### 7.3 Explicit cellular model of CRISPR gene drive

Previously, we considered a model which abstracts the mechanism of drive within cells and only deals with inheritance probabilities, *P_AB,C_*, and fitness values *f_AB_*. These were allowed to be arbitrary in our analytical calculations. However, to perform numerical simulations we must choose values for these parameters. To motivate these choices, we now formulate a model which explicitly describes how CRISPR gene drive acts within individuals.

First, we consider fitness. We assume that the wild-type has the maximum fitness of *f_s_0_s_0__* = 1 and that cost-free resistant alleles *S_i_* are identical to the wild-type allele with respect to fitness. We assume that the cost conferred by the drive is dominant (we call this cost *c*), while the cost conferred by costly resistant alleles—which are disrupted copies of the target gene—is recessive (we call this cost *s*). Furthermore, we assume—for our proposed construct—that the drive allele contains a functional copy of the target gene, so drive homozygotes do not incur the recessive cost for target disruption. Thus, we have *f_DD_* = *f_DR_i__* = *f_DS_i__* = 1 − *c*, *f_R_i_R_j__* = 1 − *s*, *f_R_i_S_j__* = *f_S_i_S_j__* = 1.

For the previously demonstrated CRISPR gene drive constructs described in the main text (Figs. 1 and 2), we again assume that disruption of the target gene produces a recessive fitness cost, s, and that the gene drive construct produces a dominant fitness cost, *c*. However, since the previously demonstrated drive constructs copied themselves by inserting at (and thus disrupting) the target sequence, we assume that the drive allele contains a disrupted copy of the target gene. Thus DD and DR individuals incur both the cost of the drive construct, *c*, and the recessive cost of resistance, *s*. We assume that these two costs are independent and thus the corresponding fitness effects are multiplicative, i.e., (1 − *c*)(1 − *s*). Thus, we have *f_DD_* = *f_DR_* = (1 − *c*)(1 − *s*), *f_DS_* = 1 − *c*, *f_RR_* =1 − *s*, *f_RS_* = *f_SS_* = 1.

Now, we consider the drive-heterozygote gamete production probabilities *p_DA_,_B_*. We assume that the proposed drive construct employs *n* gRNAs, while the previously demonstrated drive construct employs a single (*n* = 1) gRNA. Finally, we assume the cellular-level process described in the main text (Fig. 2), and this leads to the following:

- *DR_i_* individuals produce *R_i_* gametes precisely when no cutting occurs. Each of the *n* − *i* sites is susceptible to cutting, and cutting occurs independently at each with probability *q*, so we have 
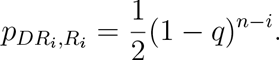
- *DR_i_* individuals produce *R_k_* gametes (with *k* > *i*) by cutting at *k* − *i* sites (where each cut occurs with probability *q*), followed by NHEJ repair (with probability 1 − *P*). Since we assume that costly resistant alleles cannot convert back to cost-free alleles, we do not consider the efficacy of repair by NHEJ. In this case, we have 
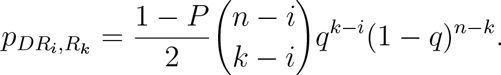
- *DR_i_* individuals produce *D* gametes by inheriting the existing *D* allele, or by cutting at one or more sites on the *R_i_* chromosome (each with probability *q*) and undergoing HR repair (with probability *P*). We have 
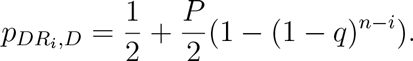
- *DS_i_* individuals produce *S_i_* gametes precisely when no cutting occurs. Each of the *n* − *i* sites is susceptible to cutting, and cutting occurs independently at each with probability *q*, so we have 
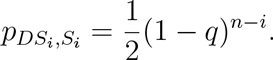
- *DS_i_* individuals produce *S_k_* gametes (with *k* > *i*) by cutting at *k* − *i* targets (each with probability q), undergoing NHEJ repair (with probability 1 − *P*), and repairing every cut perfectly (each with probability γ). We have 
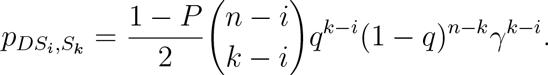
- *DS_i_* individuals produce *R_k_* gametes (with *k* > *i*) by cutting at *k* − *i* targets (each with probability *q*), undergoing NHEJ repair (with probability 1 − *P*), and repairing at least one cut imperfectly (which occurs with probability 1 − *γ^*k*−*i*^*). We have 
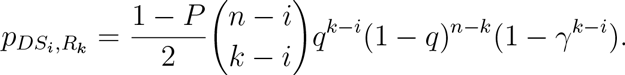
- *DS_i_* individuals produce *D* gametes by inheriting the existing *D* allele, or by cutting at one or more sites (each with probability *q*) and undergoing HR repair (with probability *P*). We have 
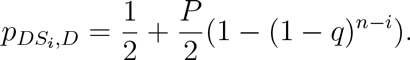

We then employ these values in the numerical simulations shown in the main text (Figs. 2C and 3), using *q* = *P* = 0.95 and γ = 1/3, with *n* = 5 for the proposed construct and *n* = 1 for the previous constructs.

## References

1. A. Burt, Site-specific selfish genes as tools for the control and genetic engineering of natural populations. Proc. Biol. Sci. 270, 921–928 (2003).

2. K. M. Esvelt, a. L. Smidler, F. Catteruccia, G. M. Church, Concerning RNA-guided gene drives for the alteration of wild populations. Elife. 3, e03401 (2014).

3. B. O. S. Akbari et al., Safeguarding gene drive experiments in the laboratory. Science. 349, 927–9 (2015).

4. K. A. Oye et al., Regulating gene drives. Science. 345, 626–8 (2014).

5. V. M. Gantz et al., Highly efficient Cas9-mediated gene drive for population modification of the malaria vector mosquito Anopheles stephensi. Proc. Natl. Acad. Sci., 201521077 (2015).

6. A. Hammond et al., A CRISPR-Cas9 gene drive system targeting female reproduction in the malaria mosquito vector Anopheles gambiae. Nat. Biotechnol. (2015), doi:10.1038/nbt.3439.

7. J. E. DiCarlo, A. Chavez, S. L. Dietz, K. M. Esvelt, G. M. Church, Safeguarding CRISPR-Cas9 gene drives in yeast. Nat. Biotechnol. (2015), doi:10.1038/nbt.3412.

8. V. M. Gantz, E. Bier, The mutagenic chain reaction: A method for converting heterozygous to homozygous mutations. Science. 348, 442–444 (2015).

9. S. P. Sinkins, F. Gould, Gene drive systems for insect disease vectors. Nat. Rev. Genet. 7, 427–435 (2006).

10. N. Windbichler et al., A synthetic homing endonuclease-based gene drive system in the human malaria mosquito. Nature. 473, 212–215 (2011).

11. O. S. Akbari et al., A synthetic gene drive system for local, reversible modification and suppression of insect populations. Curr. Biol. 23, 671–7 (2013).

12. L. Alphey, Genetic control of mosquitoes. Annu. Rev. Entomol. 59, 205–24 (2014).

13. B. Charlesworth, C. H. Langley, The population genetics of Drosophila transposable elements. Annu. Rev. Genet. 23, 251–287 (1989).

14. C.-H. Chen et al., A Synthetic Maternal-Effect Selfish Genetic Element Drives Population Replacement in Drosophila. Science. 316, 597–600 (2007).

15. C. M. Ward et al., Medea selfish genetic elements as tools for altering traits of wild populations: a theoretical analysis. Evolution. 65, 1149–62 (2011).

16. T. W. Lyttle, Segregation distorters. Annu. Rev. Genet. 25, 511–557 (1991).

17. B. Charlesworth, D. L. Hartl, Population dynamics of the segregation distorter polymorphism of drosophila melanogaster. Genetics. 89, 171–192 (1978).

18. Y. Tao, D. L. Hartl, C. C. Laurie, Sex-ratio segregation distortion associated with reproductive isolation in Drosophila. Proc. Natl. Acad. Sci. U. S. A. 98, 13183–8 (2001).

19. S. Henikoff, K. Ahmad, H. S. Malik, The centromere paradox: stable inheritance with rapidly evolving DNA. Science. 293, 1098–102 (2001).

20. A. Deredec, H. C. J. Godfray, A. Burt, Requirements for effective malaria control with homing endonuclease genes. Proc. Natl. Acad. Sci. U. S. A. 108, E874–80 (2011).

21. A. Deredec, A. Burt, H. C. J. Godfray, The population genetics of using homing endonuclease genes in vector and pest management. Genetics. 179, 2013–2026 (2008).

22. N. Windbichler et al., Homing endonuclease mediated gene targeting in Anopheles gambiae cells and embryos. Nucleic Acids Res. 35, 5922–33 (2007).

23. M. Jinek et al., A programmable dual-RNA-guided DNA endonuclease in adaptive bacterial immunity. Science. 337, 816–21 (2012).

24. P. Mali et al., RNA-guided human genome engineering via Cas9. Science. 339, 823–6 (2013).

25. L. Cong et al., Multiplex genome engineering using CRISPR/Cas systems. Science. 339, 819–23 (2013).

26. P. Mali, K. M. Esvelt, G. M. Church, Cas9 as a versatile tool for engineering biology. Nat. Methods. 10, 957–63 (2013).

27. J. A. Doudna, E. Charpentier, The new frontier of genome engineering with CRISPR-Cas9. Science. 346, 1258096–1258096 (2014).

28. M. J. Lajoie et al., Probing the limits of genetic recoding in essential genes. Science. 342, 361–3 (2013).

29. M. J. Lajoie et al., Genomically recoded organisms expand biological functions. Science. 342, 357–60 (2013).

30. M. A. Nowak, Evolutionary Dynamics (Harvard University Press, 2006).

